# Oncogenic functions of the m6A demethylase FTO in breast cancer cells involving translational upregulation of C/EBPβ-LIP

**DOI:** 10.1101/2023.09.21.558784

**Authors:** Hidde R. Zuidhof, Christine Müller, Gertrud Kortman, René Wardenaar, Ekaterina Stepanova, Fabricio Loayza-Puch, Cornelis F. Calkhoven

## Abstract

N6-methyladenosine (m6A) is a prevalent posttranscriptional mRNA modification involved in the regulation of transcript turnover, translation, and other aspects of RNA fate. The modification is mediated by multicomponent methyltransferase complexes (so-called writers) and is reversed through the action of the m6A-demethylases fat mass and obesity-associated (FTO) and alkB homolog 5 (ALKBH5) (so-called erasers). FTO promotes cell proliferation, colony formation and metastasis in models of triple-negative breast cancer (TNBC). However, little is known about genome-wide or specific downstream regulation by FTO. Here, we examined changes in the genome-wide transcriptome and translatome following FTO-knockdown in TNBC cells. Unexpectedly, FTO knockdown had a limited effect on the translatome, while transcriptome analysis revealed that genes related to extracellular matrix (ECM) and epithelial-mesenchymal transition (EMT) are being regulated through yet unidentified mechanisms. Differential translation of the CEBPB mRNA into the C/EBPβ transcription factor isoform C/EBPβ-LIP is known to act pro-oncogenic in TNBC cells through regulation of EMT genes. Here we show that FTO is required for efficient C/EBPβ-LIP expression, suggesting that FTO has oncogenic functions through regulation of C/EBPβ-LIP.

## Introduction

*N*^6^-Methyladenosine (m6A) is the most prevalent modification of mammalian mRNA and is involved in multiple steps of gene regulation (1, 2). m6A modification is performed by a methyltransferase complex consisting of methyltransferase-like 3 (METTL3), METTL14 and multiple accessory proteins including Wilms’ tumor 1-associating protein (WTAP), and is most often found in the 3’ untranslated region (3’UTR) and tends to occur at RRACH (R = G or A; H = A, C or U) consensus motifs (3-7). m6A modification may destabilize the mRNA through binding of YTH domain containing family proteins 1-3 (YTHDF1-3) (8, 9), or if found in the coding region may regulate mRNA translation (10-12). However, the transcriptome-wide effects of m6A on translation and the involvement of YTHDF1 are disputed (13, 14).

Two mammalian demethylases are known: fat mass and obesity-associated (FTO) and alkB homolog 5 (ALKBH5) (15, 16). Both are ubiquitously expressed but their targets and biological functions appear to differ. *Alkbh5*-knockout mice are anatomically normal but less fertile due to erroneous splicing and rapid degradation of genes involved in spermatogenesis (16, 17). *Fto*-knockout mice on the other hand are smaller with less lean mass and adipose tissue while no reduction in fertility has been reported (18). In humans, single nucleotide polymorphisms (SNPs) in the *FTO* gene are the strongest genetic link with obesity in genome-wide association studies (GWAS) (19). More recently, FTO was shown to play a role in many types of cancer, both with oncogenic and tumor suppressive functions depending on tumor type and/or stage (20). Mostly, FTO was found to act as an oncogene, for example in acute myeloid leukemia (AML) subtypes (21, 22), breast cancer (23) and glioma (24).

Triple-negative breast cancer (TNBC) encompasses 10-15% of all breast tumors and is characterized by loss of estrogen and progesterone receptor expression and a lack of HER2 amplification, which are therefore not available as therapeutic targets (25). TNBC has a poor prognosis with an overall 5-year relapse of approximately 40% due to metastatic spread (26). To be able to migrate to distant sites tumor cells have to undergo epithelial to mesenchymal transition (EMT) (27). Cell migration processes are regulated by genes involved in extracellular matrix modulation, cell adhesion regulation and actin cytoskeleton mobilization (28).

In tumor samples of breast cancer patients, high expression of the C/EBPβ transcription factor protein isoform C/EBPβ-LIP correlates with loss of estrogen/progesterone receptor expression, a high Ki-67 proliferative index, and lack of differentiation (29-31). Previously, we showed that C/EBPβ-LIP induces cancer-type metabolic reprogramming through the induction of aerobic glycolysis to sustain cell growth (32) and promotes EMT gene expression as well as migration and invasion of breast cancer cells and untransformed epithelial cells (33). Three C/EBPβ protein isoforms are expressed from the CEBPB mRNA through differential translation. The protein isoforms designated C/EBPβ-LAP1 and C/EBPβ-LAP2 (liver-enriched activating Proteins) are expressed through regular translation. These are transcriptional activators containing N-terminal transcriptional transactivation domains and a C-terminal basic DNA-binding and leucine-zipper dimerization domain (bZIP) (31, 34). The expression of C/EBPβ-LIP (liver-enriched inhibitory protein) is strictly dependent on the initial translation of a *cis*-regulatory small upstream open reading frame (uORF) and successive translation reinitiation from a downstream translation initiation site (for a detailed explanation, see (31)). The C/EBPβ-LIP isoform lacks the transactivation domains but does bind to DNA through the bZIP domain (35). C/EBPβ-LIP translation is stimulated by the nutrient and energy sensing mTORC1-4E-BP-eIF4E pathway, while its expression is restricted when mTORC1 signaling is suppressed and/or when under stress conditions activation of eIF-2α-kinases prevents reloading of post-uORF ribosomes with initiator Met-tRNA_i_^Met^ (31).

The molecular mechanisms and downstream targets of FTO’s function in breast cancer growth and migration remain largely unexplored. We hypothesized that FTO may contribute to these processes through regulation of general mRNA translation and/or specific mRNA translation, the latter exemplified by translation of CEBPB mRNA. To reveal the genome-wide effects of FTO expression in TNBC, we performed transcriptome and ribosome profiling experiments. While FTO knockdown hardly affected the translatome, transcriptome analysis revealed the regulation of several genes involved in extracellular matrix (ECM) and epithelial-mesenchymal transition (EMT). Similar results were obtained by treatment with the catechol-*O*-methyltransferase and the FTO inhibitor entacapone. In addition, our data show that FTO-knockdown affects C/EBPβ-LIP expression, decreasing the C/EBPβ-LIP/LAP ratio, while knockdown of the m6A-methyltransferase complex component WTAP increases C/EBPβ-LIP expression. Furthermore, we show that in TNBC cells ectopic expression of LIP rescues the migration defect caused by FTO knockdown, suggesting that LIP is involved in FTO-mediated cell migration.

## Results

### FTO facilitates breast cancer cell proliferation and migration

We examined the effect of FTO knockdown on the proliferation and migration of the breast cancer cell lines MDA-MB-231 (triple-negative breast cancer) and MCF-7 (luminal A type metastatic adenocarcinoma) using short hairpin lentiviral vectors (shFTO). FTO knockdown resulted in the expected general increase in the m6A loading of mRNAs, as measured by m6A-mRNA dot blotting **(Supplementary Figure 1A)**, while it did not affect the expression of the other known m6A demethylase ALKBH5 **(Supplementary Figure 1B)**. In a clonogenic assay, cell proliferation was significantly reduced upon FTO knockout in MDA-MB-231 and MCF-7 cells **(Figure 1A)**. The effect of FTO-knockdown on the migration of MDA-MB-231 cells was measured using a Boyden chamber (Transwell migration) assay **(Figure 1B)** and by live-cell imaging scratch wound assays using the IncuCyte system (Sartorius) **(Figure 1C)**. Both assays showed reduced cell migration upon FTO knockdown. Moreover, stable re-expression of FTO in FTO-knockdown MDA-MB-231 cells resulted in rescue of the Transwell migration **(Figure 1D)**. In addition, treatment with the FTO inhibitor entacapone caused a dose-dependent decrease in cell proliferation in both MDA-MB-231 and MCF-7 cells (**Figure 1E**) and caused a reduction in MDA-MB-231 Transwell migration (Boyden chamber) (**Figure 1F**). Together, these data and data from another laboratory (23) show that the m6A-demethylase FTO promotes proliferation and migration in breast cancer cells.

**Figure 1.**
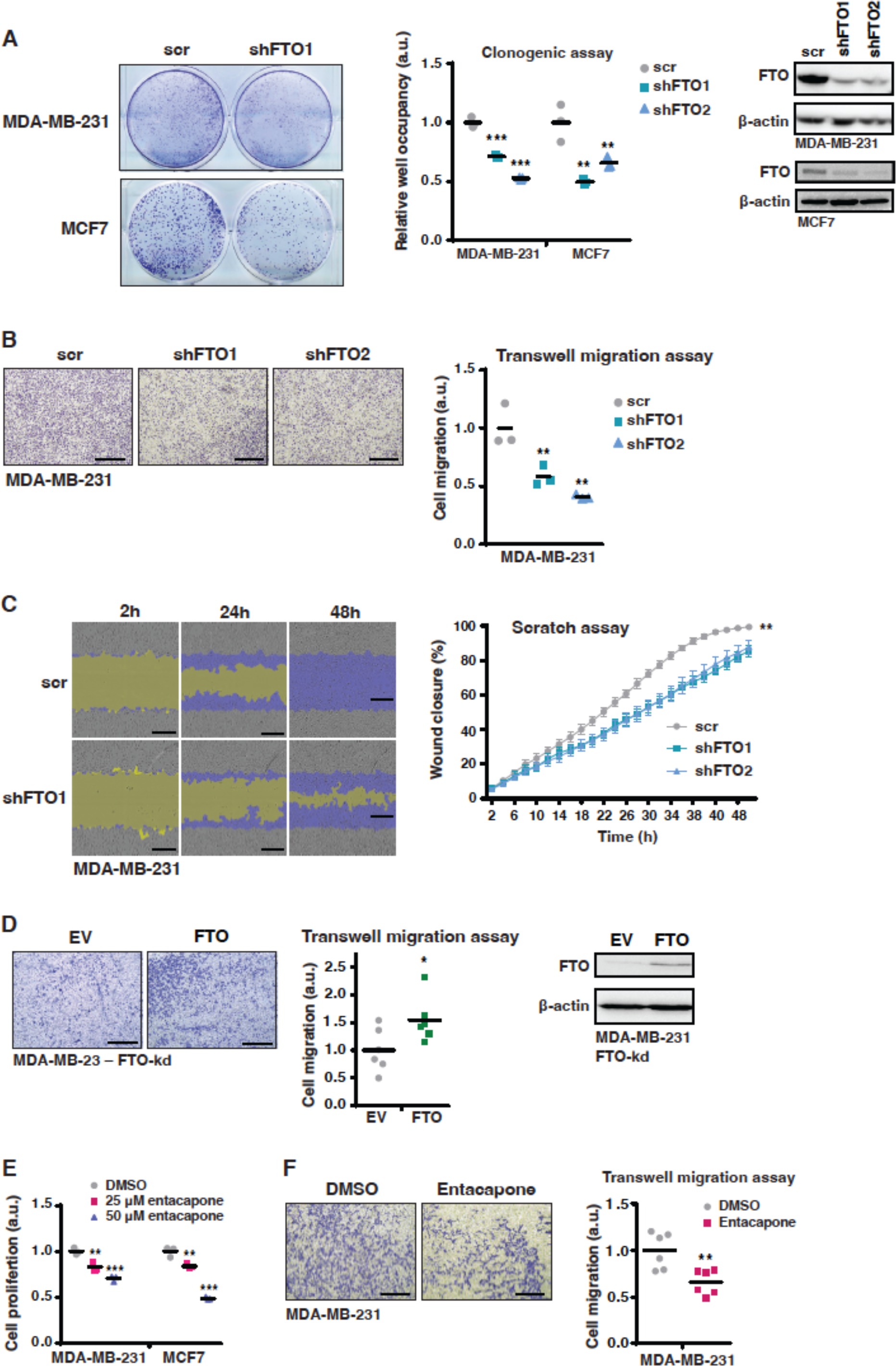
FTO knockdown impairs breast cancer cell proliferation and migration. **A**) Clonogenic assay of MDA-MB-231 and MCF-7 cells with FTO knockdown (shFTO1) or scrambled control (scr). Cells were harvested and stained after 7 days in cell culture. Quantification for two independent knockdown constructs (shFTO1 and shFTO2) is shown in the middle. At the right immunoblots showing FTO expression levels in FTO knockdown cells and scr control, and β-actin as loading control. Significance was determined by one way ANOVA with Dunnett’s posthoc test, **: p<0.01, ***: p < 0.001 (n=3 biological replicates). **B**) Transwell migration assay of MDA-MB-231 cells with FTO knockdown (shFTO1) or scrambled control (scr) at 48 hours. Scale bar represents 500 μm. Quantification is shown at the right. Significance determined by one way ANOVA with Dunnett’s posthoc test, **: p<0.01 (n=3 biological replicates). **C**) Time course of live-cell imaging scratch wound assays using IncuCyte (Sartorius) of MDA-MB-231 cells with FTO knockdown (shFTO1) or scrambled control (scr). Scale bar represents 300 μm. Quantification is shown at the right. Significance was determined by one way ANOVA with Dunnett’s posthoc test, **: p<0.01 (mean ± S.E.M., n=4 biological replicates). **D**) Transwell migration assay of MDA-MB-231 cells with FTO knockdown (shRNA1) with re-expression of FTO or empty vector (EV) control at 48 hours. Scale bar represents 500 μm. Quantification of shFTO1 and shFTO2 knockdown is shown, significance was determined by Student’s T-test, *: p<0,05, (n= 3 biological replicates). At the right immunoblots showing FTO re-expression or empty vector (EV) control in FTO knockout MDA-MB-231 cells, and β-actin as loading control. **E**) Cell viability assay of MDA-MB-231 and MCF-7 cells treated with 25 or 50 μM entacapone or DMSO control for 3 days. Significance was determined by one way ANOVA with Dunnett’s posthoc test, **: p<0.01, ***: p<0.001 (n=3 biological replicates). **F**) Transwell migration assay at 48h of MDA-MB-231 cells treated with 25 μM entacapone or DMSO. Scale bar represents 500 μm. Quantification is shown at the right. Significance determined by Student’s T-test. **: p<0.01 (n=3 biological replicates).

**Supplementary Figure 1.**
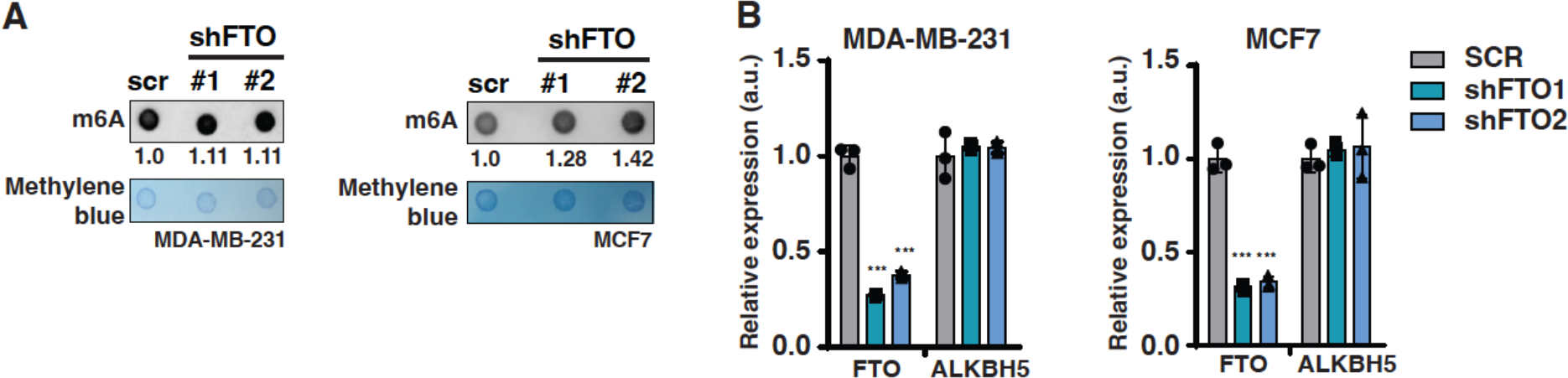
FTO knockdown impairs breast cancer cell proliferation and migration. **A**) m6A dot blot of 100ng mRNA from MDA-MB-231 cells or 150ng mRNA from MCF-7 cells with FTO knockdown (shFTO1 or shFTO2) versus scrambled-shRNA (scr) control. The blots shown are representative of 3 biological replicates used for quantification of m6A signal relative to scr control shown beneath the blot. Methylene blue staining is shown as loading control. **B**) RT-qPCR analysis of FTO and ALKBH5 gene expression in MDA-MB-231 or MCF-7 cells with FTO knockdown (shFTO1 or shFTO2) versus scrambled-shRNA control (scr). Values are normalized to scr control per gene. Significance was determined by one-way ANOVA with Dunnett’s posthoc test, ***: p < 0.001. (mean ± S.D., n=3 biological replicates).

### FTO regulates ECM and EMT gene expression

Since FTO was shown to regulate mRNA-stability we examined FTO knockdown-induced alterations in the genome-wide transcriptome. Transcriptome analysis of MDA-MB-231 cells identified 228 up- and 525 downregulated genes upon FTO knockdown (FDR < 0.05, fold change >1.5) (**Figure 2A**). The finding that more genes were downregulated than upregulated agrees with the described mRNA destabilizing effect of m6A modification (9). Gene Set Enrichment Analysis (GSEA) (https://www.gsea-msigdb.org/gsea, Broad Institute) (36) revealed a significant negative enrichment of the epithelial to mesenchymal transition (EMT) signature in FTO knockdown cells compared to control cells (**Figure 2B**). Furthermore, most of the top 15 significantly enriched gene ontology (GO) terms from the downregulated genes are related to the extracellular matrix (ECM) or EMT (**Figure 2C**). qPCR analysis confirmed the downregulation of a subset of genes by FTO knockdown, including Collagen Type XII Alpha 1 Chain (*COL12A1*), Fibronectin 1 (*FN1*), Matrix Metalloproteinase 1 (*MMP1*), Leucine Rich Repeat Containing 15 (*LRRC15*) and Tenascin C (*TNC*) (**Figure 2D**). Re-expression of FTO in FTO-knockdown MDA-MB-231 cells enhanced the expression of LRRC15 to a statistically significant level (**Figure 2E**). Overall, the results concur with the rescue of the Transwell migration phenotype upon FTO re-expression in FTO-knockdown MDA-MB-231 cells (**Figure 1D**).

**Figure 2.**
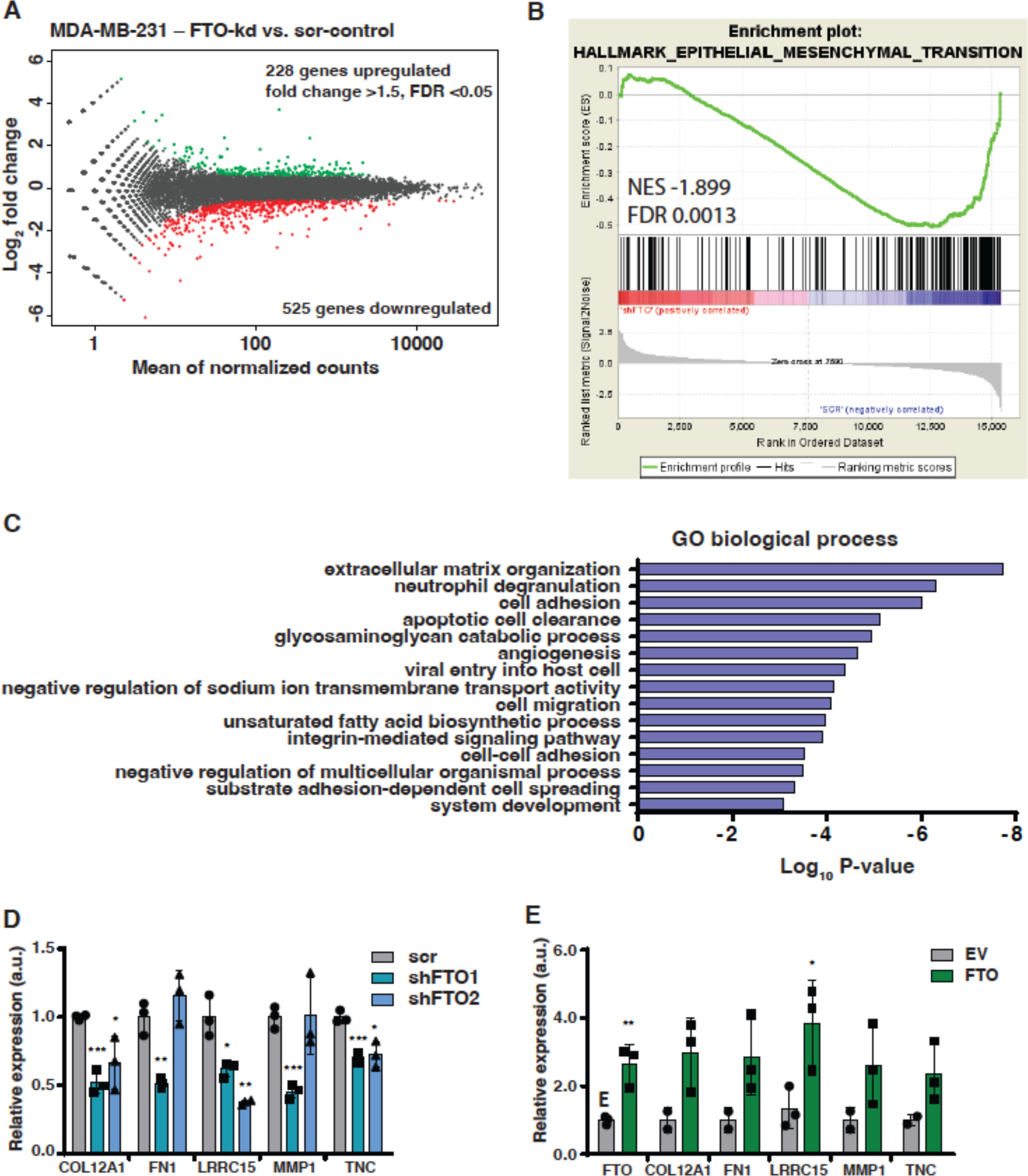
Transcriptome analysis revealing FTO-regulated ECM and EMT genes. **A**) Transcriptome analysis of MDA-MB-231 cells with FTO knockdown versus scrambled-shRNA (scr) control. Significantly regulated genes are displayed in red (fold change > 1.5; FDR<0.05). **B**) Gene set enrichment analysis (GSEA) for the hallmark epithelial mesenchymal transition shows significant negative enrichment in MDA-MB-231 FTO knockdown cells versus scrambled-shRNA (scr) control. **C**) Gene ontology (GO) term analysis of biological processes downregulated upon FTO knockdown in MDA-MB-231 cells versus scrambled-shRNA control. Significantly enriched GO-terms (p<0.001) are shown with their respective p-values (Fisher elim. topGO). **D**) RT-qPCR analysis of ECM/EMT related genes found to be downregulated upon FTO knockdown (shRNA1 or shRNA2) versus shRNA-scrambled (scr) control in MDA-MB-231 cells. Significance was determined by one way ANOVA with Dunnett’s posthoc test, *: p<0.05, **: p<0.01, ***: p < 0.001 (mean ± S.D., n=3 biological replicates). **E**) RT-qPCR analysis of ECM/EMT related genes in MDA-MB-231 FTO knockdown cells after re-expression of FTO compared to empty vector (EV) control. Significance was determined by one way ANOVA with Dunnett’s posthoc test, *: p<0.05, **: p<0.01, ***: p < 0.001 (mean ± S.D., n=3 biological replicates).

Others have screened cell signaling components for their effect on TNBC cell migration (37). Out of 4198 individual genes, 153 were identified and validated to regulate cell migration in MDA-MB-231 cells (37). We overlaid our list of downregulated genes with the validated hits from (37) and identified Low Density Lipoprotein Receptor-Related Protein 1 (*LRP1*), cAMP Responsive Element Binding Protein 5 (*CREB5*) and SWI/SNF related, matrix Associated, Actin Dependent Regulator of Chromatin, Subfamily D, Member 3 (*SMARCD3*) as the only three shared regulated genes. Upon FTO-knockdown or treatment with entacapone we observed a significant and consistent downregulation only for *SMARCD3* in MDA-MB-231 cells (**Supplementary Figure 2A**). In another study, pro-apoptotic BCL2 interacting protein 3 (BNIP3) was identified as the main gene that is downregulated by FTO through m6A demethylation. However, we failed to observe upregulation of BNIP3 in response to FTO knockdown in MDA-MB-231 cells and rather observed a trend toward downregulation of BNIP3 (**Supplementary Figure 2B**).

**Supplementary Figure 2.**
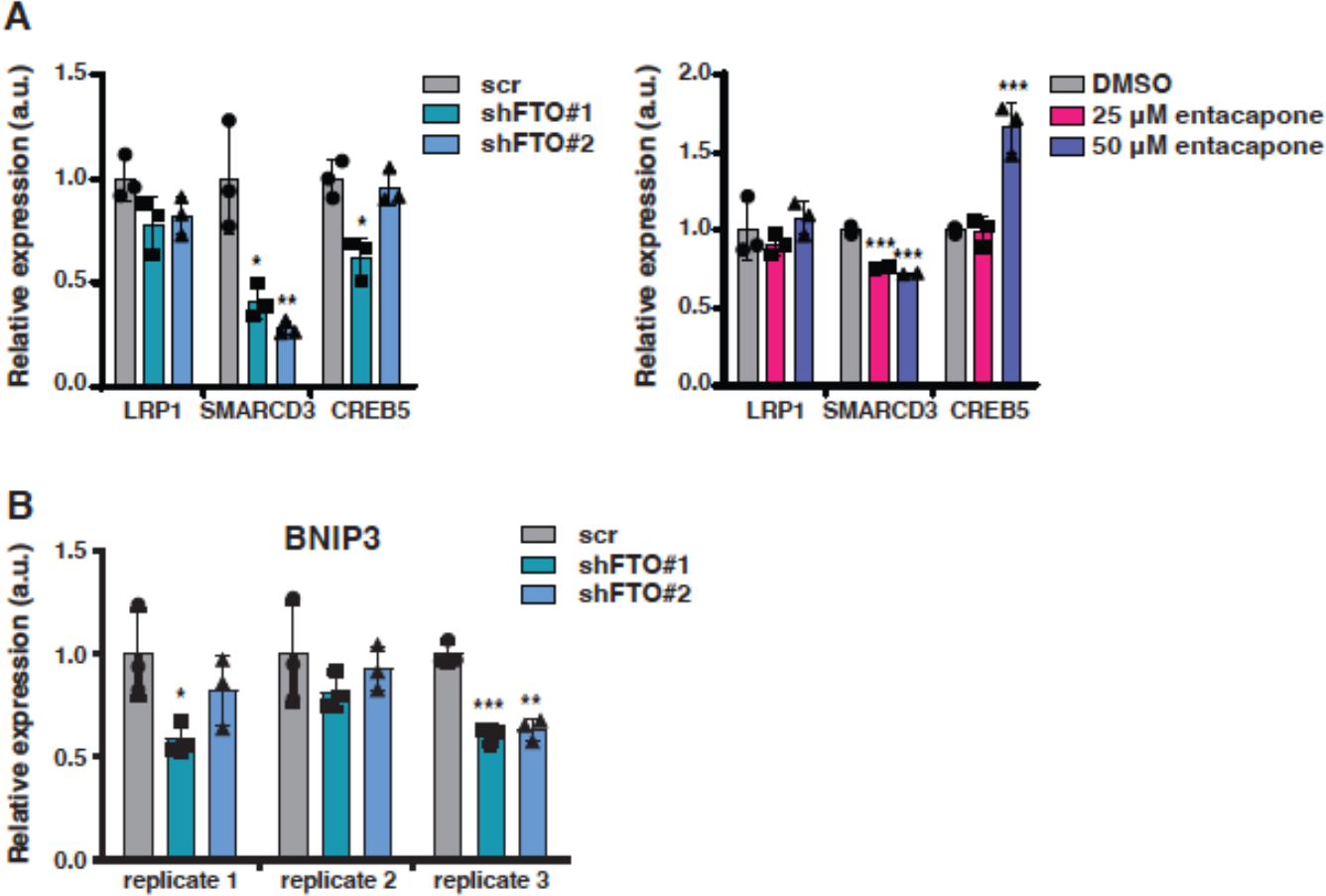
Analysis of specific genes regulation by FTO knockdown or inhibition of FTO. **A**) Left graph, RT-qPCR analysis of LRP1, SMARCD3 and CREB5 expression in MDA-MB-231 cells upon FTO knockdown (shFTO1 or shFTO2) versus shRNA-scrambled (scr) control. Significance was determined by one way ANOVA with Dunnett’s posthoc test, *: p<0.05, **: p<0.01 (mean ± S.D., n=3 biological replicates). Right graph, RT-qPCR analysis of LRP1, SMARCD3 and CREB5 expression in MDA-MB-231 cells treated with 25 or 50 μM entacapone or DMSO for 3 days. Significance was determined by one way ANOVA with Dunnett’s posthoc test, ***: p<0.001 (mean ± S.D., n=3 biological replicates). **B**) RT-qPCR analysis of BNIP3 expression in MDA-MB-231 cells with FTO knockdown (shFTO1 or shFTO2) versus shRNA-scrambled (scr) control. Significance was determined by one way ANOVA with Dunnett’s posthoc test, *: p<0.05, **: p<0.01, ***: p<0.001 (mean ± S.D., n=3 biological replicates).

### FTO does not directly regulate ECM/EMT mRNA stability

m6A-modification of mRNAs is linked to the regulation of mRNA stability by promoting degradation (9, 14). To examine whether the downregulation of ECM and EMT mRNAs in response to FTO knockdown is caused by mRNA degradation, we treated the cells with the transcription inhibitor actinomycin D and followed the degradation of selected transcripts over time. As expected, FTO mRNA levels declined in FTO-knockdown cells (**Figure 3A**). However, none of the regulated transcripts showed changes in mRNA stability upon FTO knockdown (**Figure 3B**). In addition, we did not observe an increase in mRNA-m6A levels for any of the regulated genes analyzed by methylated-RNA immunoprecipitation using a m6A antibody followed by real-time reverse transcription polymerase chain reaction (MeRIP-RT-qPCR) (**Figure 3C**). These data indicate that FTO does not regulate these transcripts directly.

**Figure 3.**
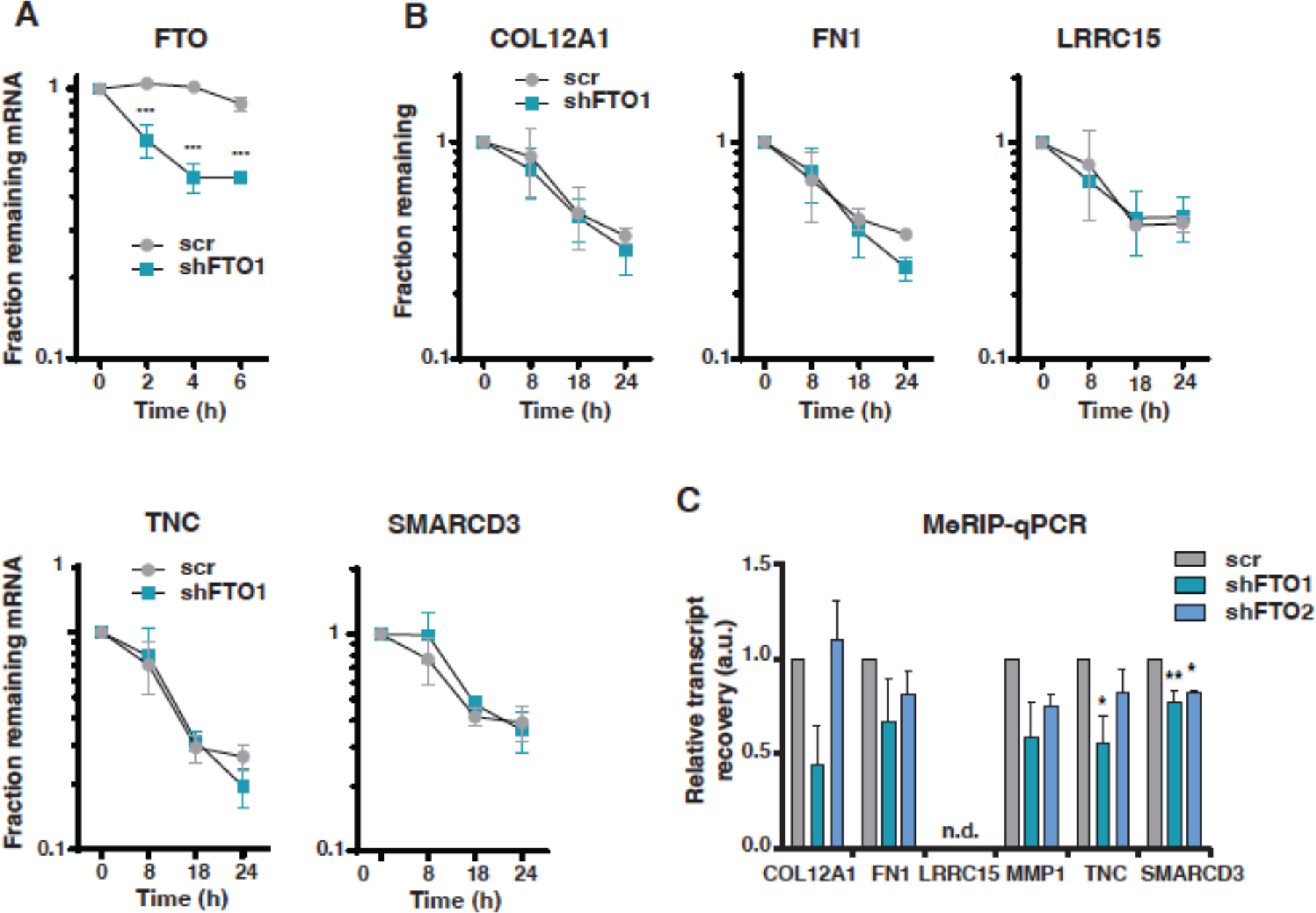
FTO does not regulate ECM/EMT target gene transcript stability or level of m6A modification. **A**) Decay curves of FTO mRNA in MDA-MB-231 cells with FTO knockdown (shFTO1) versus scrambled-shRNA (scr) control. Significance was determined by multiple Student’s T-test with Holm-Sidak correction, ***: p<0.001 (mean ± S.E.M., n=3 biological replicates). **B**) Decay curves of mRNA from indicated genes in MDA-MB-231 cells with FTO knockdown (shFTO1) versus scrambled (scr) control. Significance was determined by multiple Student’s T-test with Holm-Sidak correction (mean ± S.E.M., n=3 biological replicates). **C**) Quantification of mRNA-m6A levels of indicated genes by MeRIP-qPCR. Significance was determined by one way ANOVA with Dunnett’s posthoc test.*: p<0.05, **: p<0.01. n.d.: not detected (mean ± S.E.M., n=3 biological replicates).

### FTO deficiency does not significantly affect translation in TNBC cells

In addition to mRNA turnover, FTO function has been linked to the regulation of mRNA-translation (10, 12). To investigate FTO-dependent differences in the translatome, we performed ribosome profiling experiments with FTO knockdown and scrambled-shRNA MDA-MB-231 TNBC cells. To our surprise, the translatome was only mildly affected by FTO-deficiency and only five transcripts showed more than twofold and significantly altered translation efficiencies (FDR < 0.05, fold change >2, **Figure 4A**). The most strongly reduced in ribosome loading was DICER1, yet DICER mRNA levels determined by RT-qPCR or DICER protein levels analyzed by immunoblotting did not reveal a significant change upon FTO knockdown (**Figure 4B, C and E**). The SMAD6 and COL1A1 transcripts showed mildly reduced ribosome loading. RT-qPCR analysis showed no significant difference in expression levels (**Figure 4D**). Protein levels of COL1A1 were found reduced after long-term (20 days posttransduction) culture of FTO-knockdown cells and protein levels of SMAD6 were found lower only for the most effective shRNA1-FTO knockdown (**Figure 4E**). These analyses suggest that the FTO-mediated demethylation of the COL1A1 and SMAD6 mRNAs enhances their translation, of which FTO-COL1A1 regulation has also been observed by others (38).

**Figure 4.**
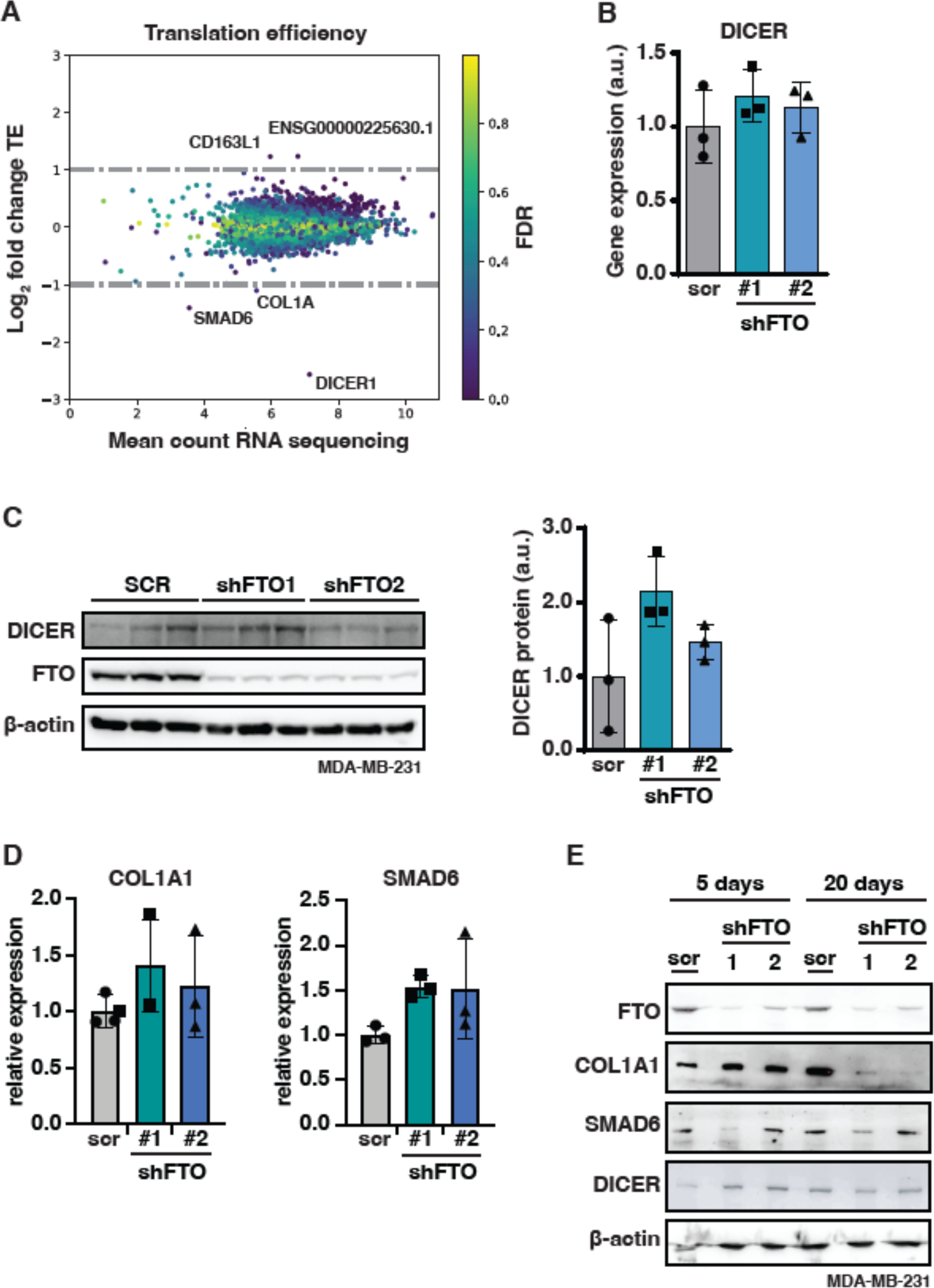
FTO knockdown has a limited effect on mRNA translation. **A**) Fold change in translation efficiency upon FTO knockdown in MDA-MB-231 cells is displayed, color coded by FDR. Significantly regulated genes above threshold are indicated by gene symbol. **B**) RT-qPCR analysis of DICER mRNA expression in MDA-MB-231 cells with FTO knockdown (shFTO1 and shFTO2) versus shRNA-scrambled (scr) control. Significance was determined by one way ANOVA with Dunnett’s posthoc test (mean ± S.E.M., n=3 biological replicates?). **C**) Immunoblots showing no change in DICER protein expression upon knockdown of FTO (shFTO) in MDA-MB-231 versus scrambled control (SCR). β-actin was used as loading control. Quantification is shown on the right. Significance was determined by one way ANOVA with Dunnett’s posthoc test (mean ± S.E.M., n=3 biological replicates?). **D**) RT-qPCR analysis of COL1A1 and SMAD6 mRNA expression in MDA-MB-231 upon FTO knockdown (shFTO1 or shFTO2) versus shRNA-scrambled (scr) control. Significance was determined by one way ANOVA with Dunnett’s posthoc test (mean ± S.D., n=3 biological replicates). **E**) Immunoblot analysis of COL1A1, SMAD6, and DICER protein expression in MDA-MB-231 upon FTO knockdown (shFTO1 or shFTO2) versus shRNA-scrambled (scr) control, and β-actin as loading control 5 days and 20 days refer to days in cell culture after transduction with shRNA retroviral vector.

The ribosome profiling experiment provides additional data about possible FTO-related amino acid or tRNA metabolism. The position of a ribosome on the mRNA-ribosome protected fragment (RPF) can be deduced from the RPF 5’ end. Using this information and the known mRNA reading frame, the codons in the ribosomal A (aminoacyl), P (peptidyl-transfer) and E (exit) sites can be determined. Positions 9, 12 and 15 from the 5’ end of the RFPs correspond to respectively the ribosomal E, P and A site, respectively. When for example a particular amino acid or tRNA is depleted from cells, this will cause an accumulation of ribosomes on the corresponding codons. Since amino acid recognition takes place at the A-site, the affected codons will be enriched at position 15 in RPFs (39). Using differential ribosome codon reading (diricore) analysis (39) no significant differences in ribosome A-site occupancy were observed (**Supplementary Figure 4B**). As no specific amino acid or tRNA shortages are induced by FTO deficiency, this indicates that FTO is not critically involved in their metabolic regulation.

**Supplementary Figure 4.**
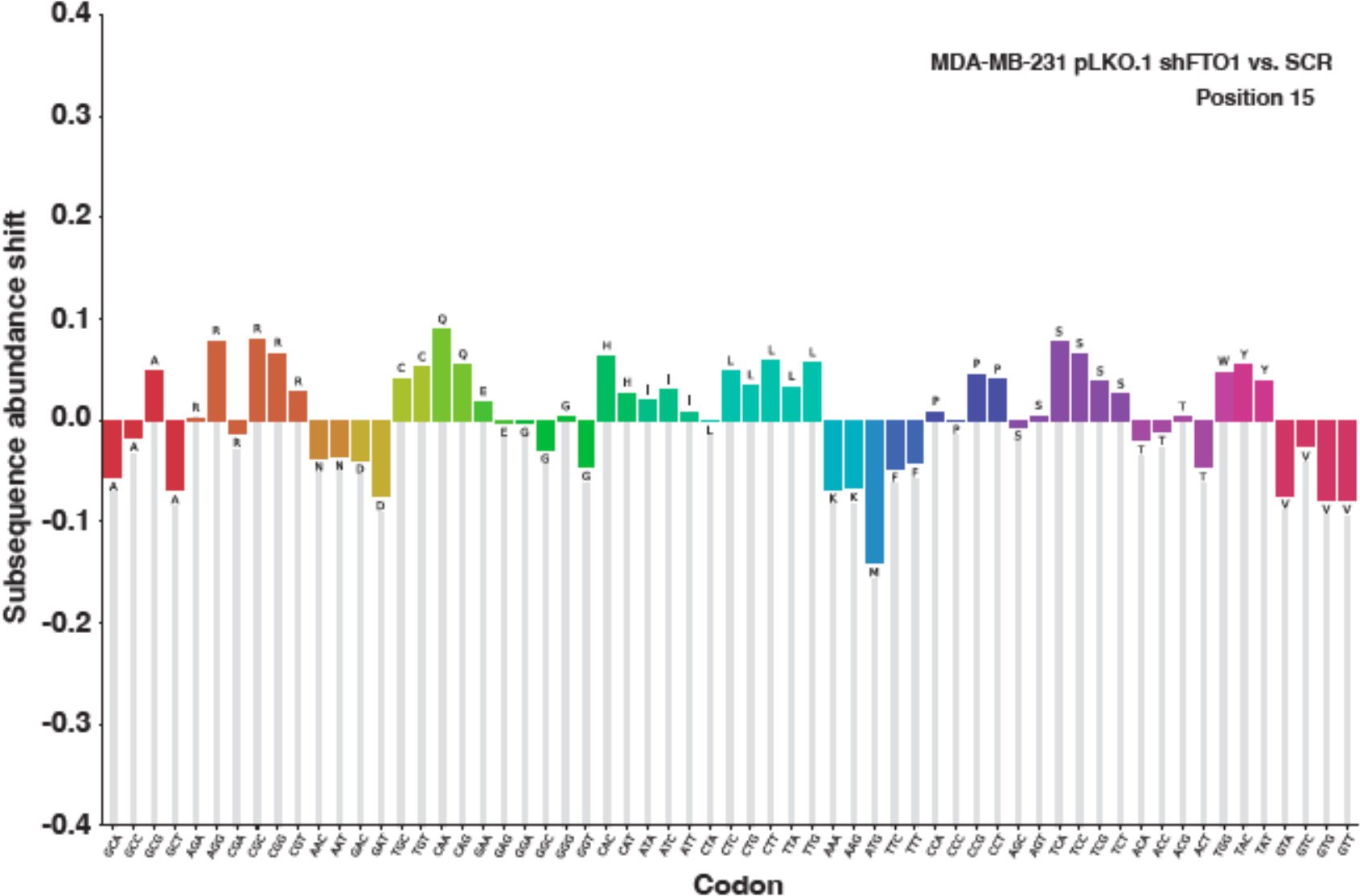
FTO knockdown does not result in specific amino acid or tRNA shortages. Subsequence abundance analysis of codons at the +15 position relative to the ribosome protected fragment (RPF) 5’-end. All transcripts in the coding frame are considered for this analysis. No significant enrichment or depletion of codons can be seen, indicating no obvious shortages of amino acids causing ribosome stalls/halts.

### FTO and WTAP regulate C/EBPβ isoform expression

The unexpectedly low number of genes that were found to be translationally regulated more than twofold raised the question whether specific translation events on single mRNAs could be affected by FTO that does not significantly alter overall ribosome loading. We chose to investigate the regulation of CEBPB mRNA that is differentially translated into three protein isoforms C/EBPβ-LAP1, -LAP2 and C/EBPβ-LIP, of which the latter involves a *cis*-regulatory upstream open reading frame (uORF)(**Figure 5A**)(31). We have shown previously that translational upregulation of C/EBPβ-LIP is involved in breast cancer cell proliferation, migration, and metabolism (32, 40). Sequence-based RNA adenosine methylation site predictor (SRAMP, http://www.cuilab.cn/sramp) identified six putative m6A sites with the highest confidence level in the CEBPB mRNA (41). Published data on m6A individual-nucleotide-resolution crosslinking and immunoprecipitation (miCLIP) experiments confirmed m6A modification of four of these sites (42), while antibody-free deamination adjacent to RNA modification targets (DART)-seq identified two additional sites (43). In concordance with the genome-wide distribution of m6A on transcripts (6), all but one of these sites are in the 3’UTR of the CEBPB mRNA **(Figure 5A)**. In a study using m^6^A-level and isoform-characterization sequencing (m^6^A-LAIC-seq), CEBPB mRNA was found to be m6A-modified in a human embryonic stem cell or B-cell lymphoblastoid cell line (44). To study whether m6A modification of the CEBPB mRNA contributes to its translational regulation, we applied FTO knockdown (shFTO) in triple-negative MDA-MB-231 and luminal A-type MCF-7 breast cancer cell lines, and mouse embryonic fibroblasts (MEFs). Knockdown of FTO decreased C/EBPβ-LIP levels with a concomitant decrease in the C/EBPβ-LIP/LAP ratio in MDA-MB-231 cells **(Figure 5B)** and MCF-7 cells **(Supplementary Figure 5A)**, and to a lesser extent in MEFs **(Supplementary Figure 5B)**. At least in MDA-MB-231 cells, FTO-knockdown did not affect CEBPB mRNA stability **(Figure 5C)**. Conversely, knockdown of WTAP, a member of the m6A-methyltransferase complex, increased the expression of C/EBPβ-LIP in MDA-MB-231 cells **(Figure 5D)**. To directly examine FTO-related changes in CEBPB mRNA-m6A levels, we performed methylated RNA immunoprecipitation using a m6A antibody followed by real-time reverse transcription polymerase chain reaction (MeRIP-RT-qPCR). Recovery after IP over input was determined as a reflection of transcript m6A levels. A clear trend toward increased CEBPB mRNA m6A-levels upon FTO knockdown was observed that did not reach statistical significance because of high experimental variation **(Figure 5E)**. Taken together, the data suggest that reversible WTAP- and FTO-controlled m6A modification of CEBPB mRNA alters its translation.

**Figure 5.**
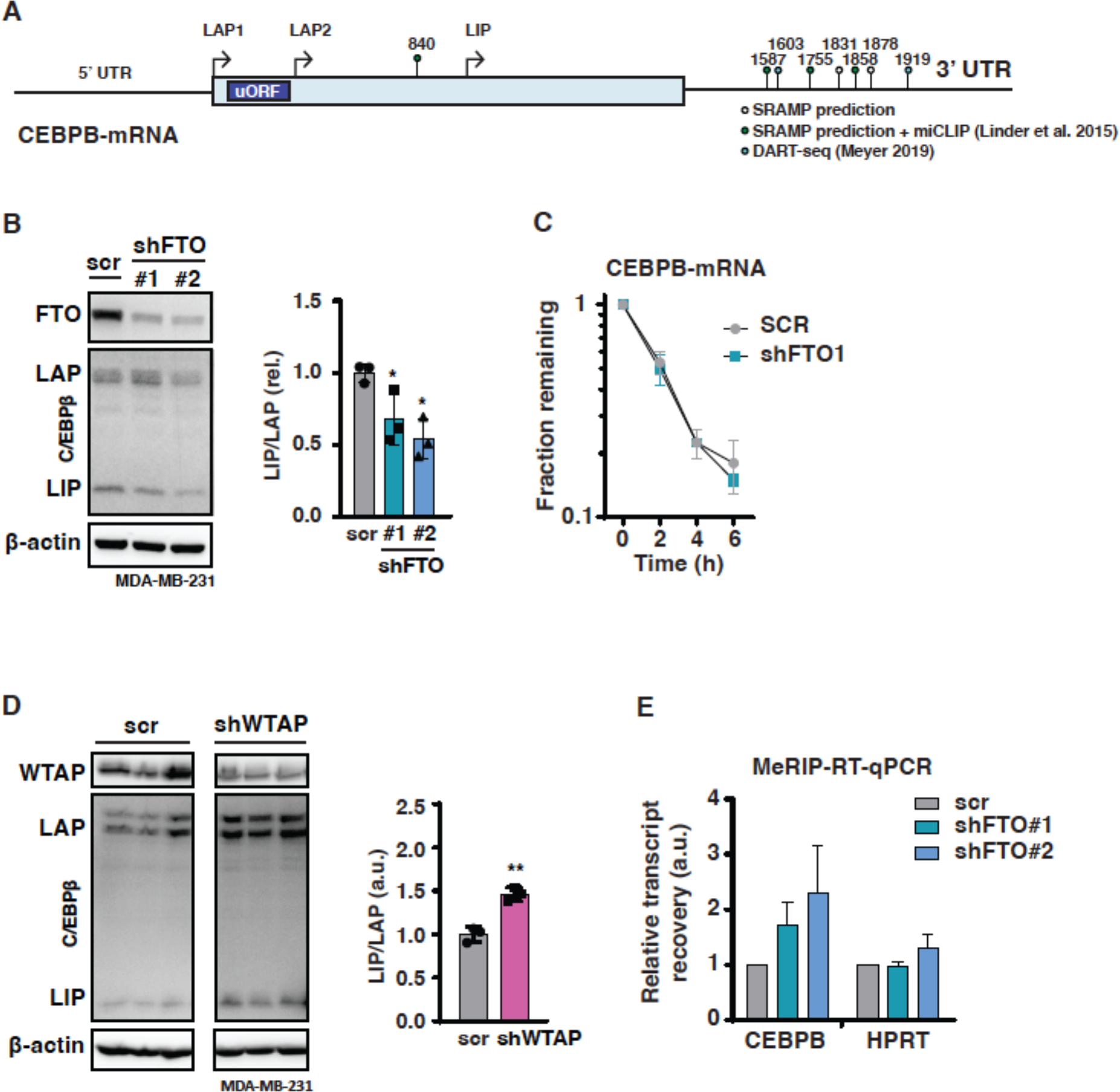
FTO or WTAP knockdown reciprocally regulates C/EBPβ isoform expression. **A**) Schematic overview of the human CEBPB mRNA with indicated translation start sites for LAP1, LAP2, LIP and the uORF. Predicted (SRAMP, (42)) and experimentally detected m6A modifications from HEK cells (miCLIP: Linder et al. 2015 (43), DARTseq Meyer 2019 (44)) are indicated with adenosine base number. **B)** Immunoblot showing reduced levels of C/EBPβ-LIP upon knockdown of FTO (shFTO) in MDA-MB-231 cells versus scrambled control (SCR). β-actin was used as loading control. **C**) Decay curves of CEBPB mRNA from MDA-MB-231 cells with FTO knockdown (shFTO1 or shFTO2) versus shRNA-scrambled (scr) control. Values represent mean ± S.E.M. of 3 biological replicates. Significance was determined by multiple Student’s T-test with Holm-Sidak correction. **D**) Immunoblot showing increased levels of C/EBPβ-LIP upon knockdown of WTAP (shWTAP) in MDA-MB-231 and BT-20 cells versus scrambled control (SCR). β-actin was used as loading control. **E**) Quantification of C/EBPβ-LIP/LAP ratio in MDA-MB-231 or BT-20 cells with WTAP knockdown. Significance was determined by Student’s T-test. **: p<0.01, ***: p<0.001 (n=3 biological replicates).

### C/EBPβ-LIP overexpression partially rescues the migration phenotype of cells with FTO knockdown

Since we showed earlier that LIP stimulates the migration of breast cancer as well as untransformed breast epithelial cells (40), we investigated whether overexpression of C/EBPβ-LIP would reverse the reduced MDA-MB-231 cell migration phenotype caused by FTO knockdown. Overexpression of C/EBPβ-LIP in wt MDA-MB-231 control cells significantly further increased cell migration (Boyden chamber) in one of the two independent experiments (**Figure 6A and B, and Supplementary Figure 6A and B)**. FTO knockdown strongly reduced cell migration in all cases, as expected. C/EBPβ-LIP overexpression efficiently rescued cell migration in shFTO2-knockout cells and moderately rescued migration in one of the shFTO1-knockout cells. Altogether, the data indicate that (m6A)-demethylation of CEBPB mRNA facilitates C/EBPβ-LIP upregulation, which could contribute to the promigratory phenotype induced by FTO.

**Figure 6.**
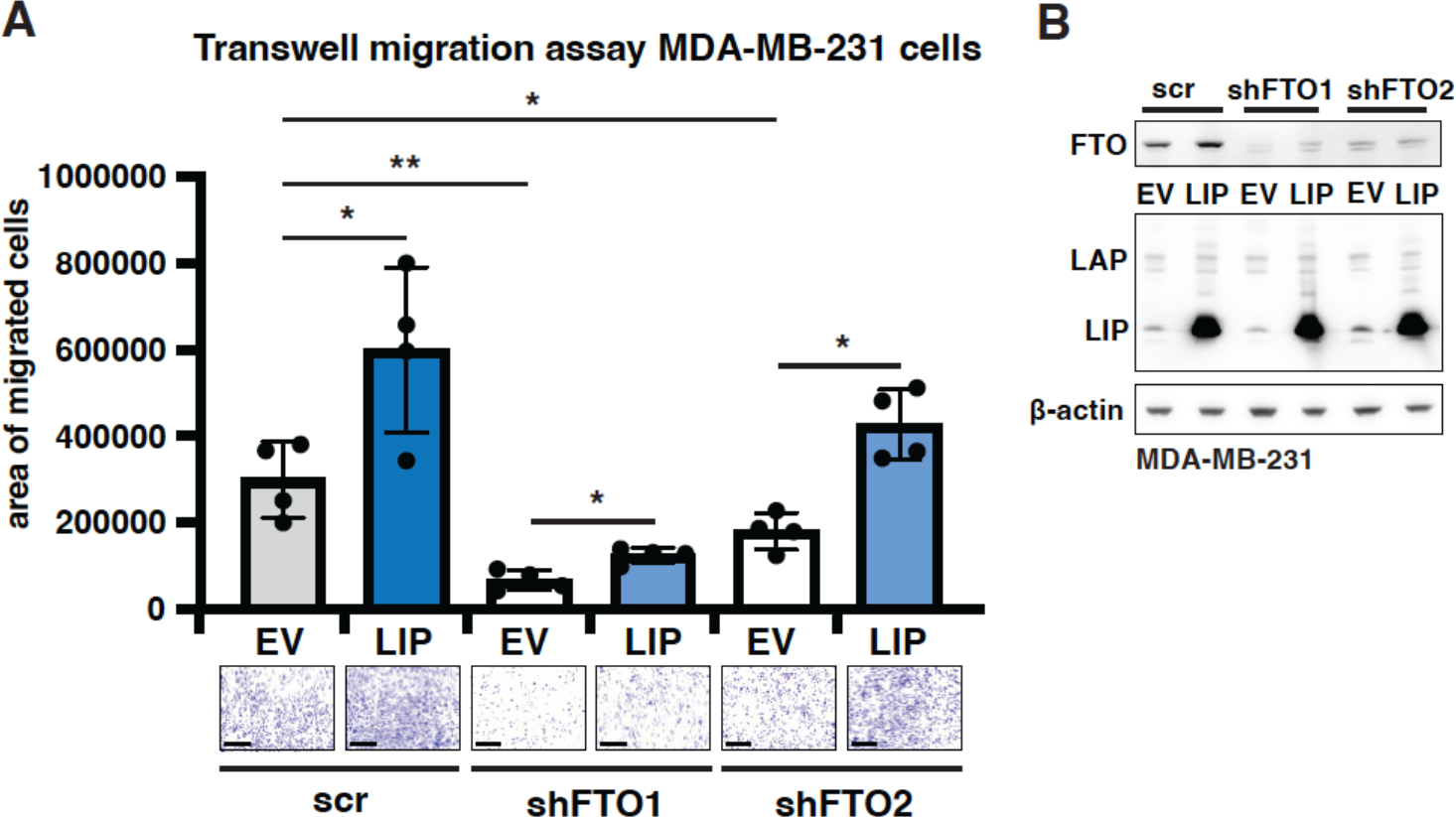
Ectopic expression of C/EBPβ-LIP enhances cell migration independent of FTO. **A**) Transwell migration assay of MDA-MB-231 cells with FTO-knockdown (shFTO1) or scrambled control (scr) with overexpression C/EBPβ-LIP or empty vector (EV) control, at 48 hours. The graph shows quantification of migrated cells with pictures of stained cells presented at the bottom. Scale bar represents 500 μm. Significance determined by one way ANOVA with Dunnett’s posthoc test, **: p<0.01 (n=4 technical replicates). **B**) Immunoblot showing overexpression of C/EBPβ-LIP in FTO knockdown (shFTO1 or shFTO2) or scrambled-shRNA (scr) control MDA-MB-231 cells. β-actin was used as loading control.

**Supplementary Figure 6.**
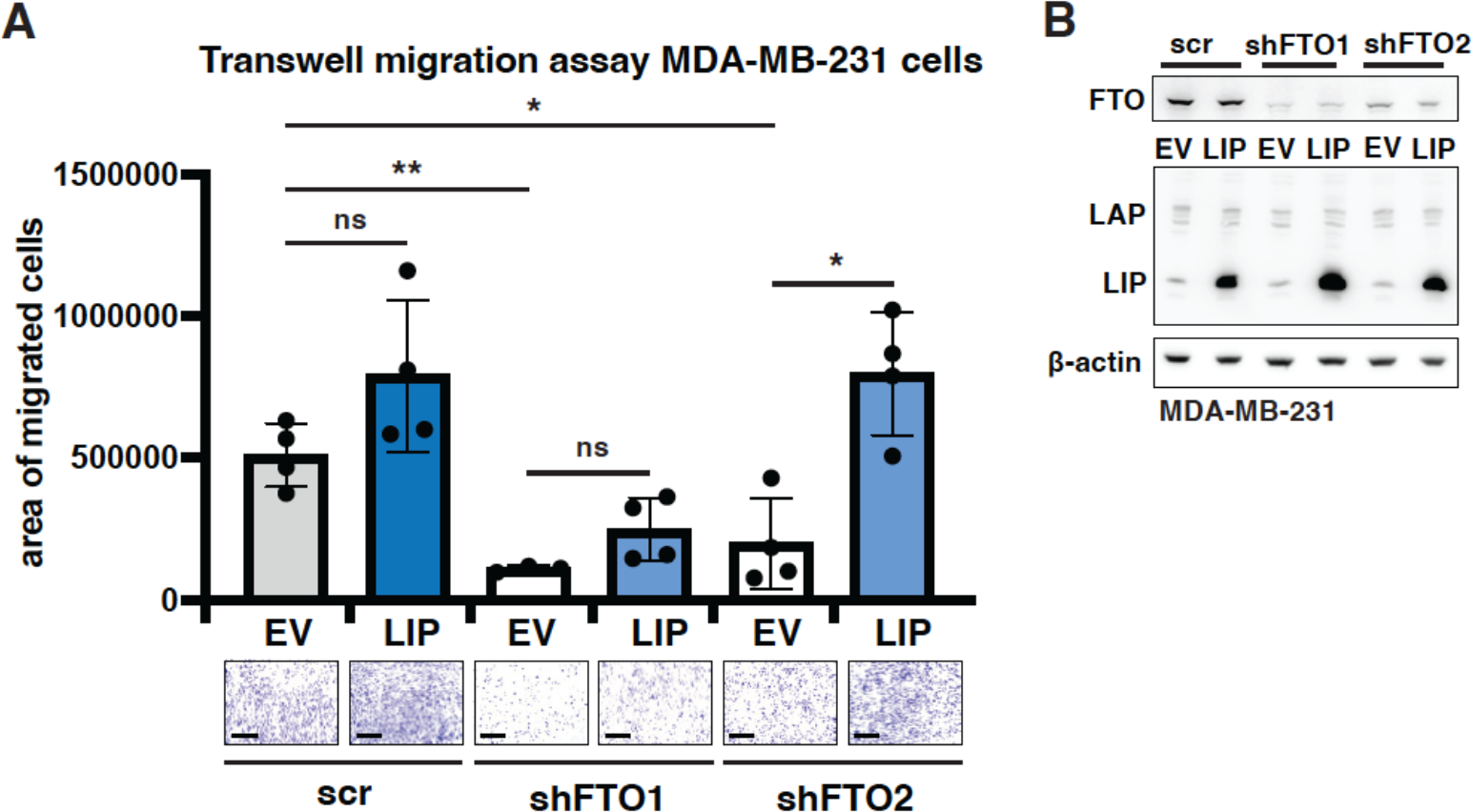
Ectopic expression of C/EBPβ-LIP enhances cell migration independent of FTO. **A**) Second transwell migration assay of MDA-MB-231 cells with FTO knockdown (shFTO1) or scrambled control (scr) with overexpression C/EBPβ-LIP or empty vector (EV) control, at 48 hours. The graph shows quantification of migrated cells with pictures of stained cells presented at the bottom. Scale bar represents 500 μm. Significance determined by one way ANOVA with Dunnett’s posthoc test, **: p<0.01 (n=4 technical replicates). **B**) Immunoblot showing overexpression of C/EBPβ-LIP in FTO-knockdown (shFTO1 or shFTO2) or scrambled-

## Discussion

Here we present data showing that FTO expression stimulates the proliferation and migration of MDA-MB-231 (triple-negative breast cancer) and MCF-7 (luminal A type metastatic adenocarcinoma) breast cancer cells in cell culture. Our transcriptome analysis and ribosome profiling experiment using MDA-MB-231 cells and FTO knockdown showed little effect on the translatome but revealed significant regulation of transcripts involved in the ECM and EMT. In addition, the data show that FTO knockdown reduces the expression of C/EBPβ-LIP, while WTAP knockdown results in higher levels of C/EBPβ-LIP, suggesting that FTO-mediated m6A demethylation is involved in the regulated translation of the CEBPB mRNA into its protein isoforms C/EBPβ-LAP1/2 and C/EBPβ-LIP.

A role of FTO-mediated m6A demethylation in the development of cancer progression has emerged in recent years. While tumor suppressive functions for FTO have been described in for example colorectal cancer and hepatocellular carcinoma, reports describing oncogenic functions in different cancer types including lung cancer, glioma and AML are far more abundant (reviewed in (20)). Analysis of broad collections of TNBC cells and samples suggests that FTO function must be maintained for TNBC cell survival. The catalog of somatic mutations in cancer (COSMIC, https://cancer.sanger.ac.uk/cosmic) database reports only a single intronic mutation in the *FTO* gene in 117 TNBC samples analyzed. Furthermore, in the 83 breast cancer cell lines listed in the cancer cell line encyclopedia (CCLE) dependency map portal (depmap, https://depmap.org/portal/) only a single cell line with an intronic mutation was found.

Our results and data presented by Niu *et al* (23) show that FTO function is required for breast cancer cell proliferation and migration. The study by Niu *et al* (23) showed that FTO is upregulated in breast tumor samples together with downregulation of METTL3/14, ALKBH5, YTHDC2 and YTHDF1, and is associated with an overall decline in global mRNA m6A levels. Furthermore, they showed that in breast cancer patients high FTO levels correlate with poorer survival. Niu *et al* (23) identified the pro-apoptotic BNIP3 (RNA degradation by FTP) as the downstream target of FTO by transcriptome and m6A-seq analysis using MDA-MB-231 FTO-knockdown and MCF-7 cells treated with the FTO inhibitor GSE3188. They also showed that BNIP3 expression alleviates FTO-dependent tumor growth in a xenograft mouse model. However, we did not observe such regulation of BNIP3 in MDA-MB-231 cells and rather observed downregulation upon FTO-knockdown.

Instead, our study shows that FTO knockdown inhibits and WTAP knockdown stimulates translation into C/EBPβ-LIP, which can act as an oncogene and regulates EMT/ECM genes in TNBC (40). This suggests that for efficient uORF-dependent regulation of LIP, FTO-mediated CEBPB mRNA demethylation is needed. We show that ectopic expression of C/EBPβ-LIP stimulates cell migration in FTO knockdown cells, suggesting that at least part of the effects of FTO on cell migration occur through regulation of C/EBPβ-LIP. The question that arises from this observation is whether FTO collectively regulates mRNAs containing uORFs or other translation *cis*-regulatory elements. In line with this hypothesis, amino acid starvation was shown to regulate the translation activating transcription factor 4 (ATF4) via m6A (45). ATF4 translation depends on reinitiation following uORF translation, allowing bypassing of an inhibitory second uORF when eIF2-GTP levels are low (46). FTO or ALKBH5 knockdown represses ATF4 reinitiation, whereas knocking down the m(6)A methyltransferase subunits METTL3 or METTL14 promotes ATF4 translation (45). Translation initiation site profiling experiments using the inhibitor harringtonine, which stalls ribosomes at the initiation sites (47), should be performed to investigate whether FTO translational regulation is more about start-site selection than general translation efficiency.

The translatome experiment only showed very limited changes upon FTO knockdown, with only two transcripts being translationally upregulated and three transcripts being translationally downregulated. Immunoblotting experiments showed that COL1A1 and SMAD6 protein levels were accordingly downregulated, yet that DICER protein levels were not changed, although DICER was the most strongly regulated mRNA. It is possible that FTO knockdown did not reach levels low enough to broadly affect translation, and FTO knockout would have been a better strategy to reveal such regulation. Nonetheless, in a (patho)physiological setting, one would rather expect levels of FTO to vary than to be lost completely. For example, in humans, a recessive loss-of-function mutation in *FTO* leads to postnatal growth retardation, microcephaly, facial dimorphism, functional brain deficits and death within 30 months of age (48).

Taking into account the many studies on the diverse and partially contradictory roles of FTO in cancer (reviewed in (20, 49)), one must conclude that the effects of FTO on specific and global regulation of RNA demethylation can only be understood and eventually employed for therapeutic strategies in specific situations, e.g., a specific cancer type at a specific stage.

## Materials & Methods

### Cell culture

MDA-MB-231 and HEK cells were cultured in DMEM, high glucose with GlutaMAX and pyruvate (Gibco) supplemented with 10% FCS, 10 mM HEPES and penicillin/streptomycin. MCF-7 cells were cultured in RPMI 1640 with GlutaMax and 25 mM HEPES (Gibco) supplemented with 10% FCS, 1 mM pyruvate and penicillin/streptomycin. All cell lines were incubated at 37 °C with 5% CO_2_ in a humidified incubator.

### DNA constructs

Short hairpin RNA sequences targeting FTO or nontarget ‘scrambled’ shRNA control (Sigma-Aldrich) were ordered as forward and reverse oligos, they were annealed and inserted into the pLKO1puro vector using AgeI and EcoRI restriction sites. ShFTO1 fw: 5’-CCG GTC ACC AAG GAG ACT GCT ATT TCT CGA GAA ATA GCA GTC TCC TTG GTG ATT TTT G-3’, shFTO1 rv: 5’-AAT TCA AAA ATC ACC AAG GAG ACT GCT ATT TCT CGA GAA ATA GCA GTC TCC TTG GTG A-3’, shFTO2 fw: 5’-CCG GTA GTC TGA CTT GGT GTT AAA TCT CGA GAT TTA ACA CCA AGT CAG ACT ATT TTT G-3’, shFTO2 rv: 5’-AAT TCA AAA ATA GTC TGA CTT GGT GTT AAA TCT CGA GAT TTA ACA CCA AGT CAG ACT A-3’, shWTAP fw: 5’-CCG GAT GGC AAG AGA TGA GTT AAT TCT CGA GAA TTA ACT CAT CTC TTG CCA TTT TTT G-3’, shWTAP rv: 5’-AAT TCA AAA AAT GGC AAG AGA TGA GTT AAT TCT CGA GAA TTA ACT CAT CTC TTG CCA T-3’. For rescue of FTO expression, human FTO (from pCMV-FTO, HG12125-UT, Sino Biological) was subcloned and inserted into the pcDNA3.1+ expression vector using KnpI/XbaI. For ectopic LIP expression, human LIP was cloned and inserted into the lentiviral expression vector pLVX-IRES-neo as described in (40).

### Knockdown cells

For knockdown cells, 3.8 x 10^6^ HEK cells were seeded in 10 cm dishes one day prior to calcium phosphate transfection of pLKO.1 constructs along with pMDL-RRE, pCMV-VSVg and pRSV-Rev expression constructs. After 2 days virus was harvested and used for transduction of target cells followed by selection with either 0.25 μg/ml (MDA-MB-231) or 0.5 μg/ml (MCF-7) puromycin. For ectopic LIP expression concomitant with nontarget ‘scrambled’ sh control RNA or FTO sh RNAs in MDA-MB-231 cells, HEK cells were cotransfected with pLKO.1 constructs and either human LIP pLVX-IRESneo vector or empty pLVX-IRESneo vector as a control along with pMDL-RRE, pCMV-VSVg and pRSV-Rev expression constructs. Virus was harvested and used for transduction of target cells as described above and cells were selected with both puromycin (0.5 μg/ml) and Geneticin (1 mg/ml). For FTO-overexpressing/rescued cells, MDA-MB-231 cells were seeded in 6-well plates one day prior to FuGene transfection with pcDNA3.1+ empty vector or FTO according to the manufacturer’s instructions. Cells were selected with 700 μg/ml Geneticin.

### Immunoblotting

Cells were washed twice in ice-cold PBS prior to harvesting by scraping in ice-cold PBS. Cells were lysed by sonication in 50 mM Tris pH 7.5, 150 mM NaCl, 1 mM EDTA, and 1% Triton-X 100 supplemented with protease and phosphatase inhibitors (Roche). Equal amounts of protein were separated by SDS-PAGE using Mini-PROTEAN® TGX™ Precast Protein Gels (#4561094, Bio-Rad) and transferred to 0.2 μM PVDF membranes by a Trans-Blot Turbo System (Bio-Rad), using RTA Midi kit (#1704273 Bio-Rad) following the manufacturer’s instructions. Blots were washed in TBS-T (50 mM Tris, 150 mM NaCl, 0,02% Tween-20) and blocked/stained in 5% milk or BSA in TBS-T. Following overnight incubation with primary antibody, blots were stained with secondary antibodies (mouse or rabbit IgG HRP-linked antibody, Thermo Fisher, 1:5000), and signals were visualized by chemiluminescence (ECL, Amersham) on an ImageQuant LAS 4000 mini biomolecular imager (GE Healthcare). The supplied software was used for image quantification. The following primary antibodies were used for detection: FTO: NB110-60935, Novus Biologicals (1:1666); β-Actin: 691001, MP Bio (1:5000); C/EBPβ: ab32358, Abcam (1:1000); WTAP: #56501 (1:1000), Cell Signaling; COL1A1: #72026 (1:1000), Cell Signaling; SMAD6: NB100-56440, Novus Biologicals (1:1000). DICER protein levels were examined by tris-acetate immunoblotting as described (50) with minor modifications. Briefly, 80 μg protein per sample was loaded on 7% Tris-acetate polyacrylamide gels (3% stacking) and transferred overnight at 20 V to 0.45 μm PVDF membrane by tank blotting in bicine/bis-tris transfer buffer. The membranes were further processed as described, and the primary antibody was DICER (sc-136979, Santa Cruz (1:1000).

### RT-qPCR

RNA was isolated from cells using the RNeasy Kit (Qiagen) according to the manufacturer’s instructions. 1 μg of total RNA was reverse transcribed using a Transcriptor First Strand cDNA Synthesis Kit (Roche) with random hexamer primers. Gene expression was analyzed using SYBR Green Master Mix (Bio-Rad) on a LightCycler 480 (Roche) and normalized to β-actin expression. Oligo sequences are listed in Supplementary Table 1.

### Cell migration assays

Cells were serum starved in their canonical medium with 0.1-1% serum. For scratch assays, 15,000 MDA-MB-231 cells were seeded in 96-well ImageLock plates compatible with the Incucyte Zoom system (Sartorius). 24 hours after seeding a scratch wound was made using the WoundMaker Tool (Sartorius) and the cells were incubated in low-serum medium after washing twice. Cells were imaged every 2 hours for 48 hours and the relative wound density was calculated using the provided software according to the manufacturer’s instructions. For Transwell migration assays, 20,000 cells were seeded in 0.1-1% serum in the upper compartment of 8 μm Corning Transwell polycarbonate membrane cell culture inserts (3422, Corning) and attracted to the bottom by medium with 10% serum. After 48 hours, the cells were stained in 96% ethanol on ice and stained with 0.5% crystal violet in 10% ethanol in water. Inserts were rinsed with demi water, air dried and imaged using a Leica DMC2900 digital microscope. For quantification, at least 3 inserts with 3 random images from each insert per condition were analyzed using ImageJ. To determine the effect of entacapone on cell migration, cells were pretreated with 25 μM entacapone (SML-0654, Sigma-Aldrich) for 24 h prior to seeding as described with 25 μM entacapone in both low serum and attractant media.

### Clonogenic and cell viability assays

For clonogenic assays, 3,000 cells were seeded in 5 ml of their normal medium in 6-well plates and incubated for 7 days. Cells were washed twice in PBS, fixed in 96% ethanol on ice and washed again twice in PBS before staining with 0.5% crystal violet in 10% ethanol in water. Wells were washed with demi water, air dried and imaged prior to quantification using ImageJ. For cell viability assays, 2,500 (MDA-MB-231) or 7,500 (MCF-7) cells were seeded per well in 96-well plates with the indicated drug concentrations of entacapone and/or HLM006474 (SML1260, Sigma-Aldrich). After 72 hours cell viability was determined using a CellTiter-Fluor™ Cell Viability Assay (Promega) according to the manufacturer’s instructions.

### m6A dot blotting

mRNA was isolated from total RNA using Dynabeads™ Oligo(dT)25 (Thermo Fisher) and diluted to 50 (MDA-MB-213) or 75 (MCF-7) ng/μl in RNase free water. Samples were denatured for 3 minutes at 95 degrees and 2 μl was spotted on Hybond®-N+ hybridization membrane (Amersham). mRNA was crosslinked to the membrane using a Stratalinker 2400 UV crosslinker (twice 120,000 μJ/cm^2^) and the membrane was washed for 10 minutes in TBS-T. Membranes were stained with 0.05% methylene blue in 0.5 M acetic acid for 15 minutes and washed three times for 2 minutes in Milli-Q water. Following imaging for loading control the membrane was blocked in 5% BSA in TBS-T and incubated overnight at 4 °C using N6-methyladenosine antibody (ab151230, Abcam). The next day, the membranes were washed three times in TBS-T, stained with 1:5000 rabbit IgG HRP-linked antibody (Thermo Fisher) for 1 hour and washed again three times in TBS-T. The m6A signal was developed using chemiluminescence as described for immunoblotting.

### Ribosome profiling

Preparation of ribosome-protected footprints for deep sequencing was essentially performed as described (51). Briefly, 30 x 10^6^ cells were harvested in ice cold PBS with 100 μg/ml cycloheximide (Sigma-Aldrich) and 10% was kept aside for RNA-sequencing. Cells were lysed in lysis buffer (20 mM Tris-HCl pH 7.5, 10 mM MgCl_2_, 100 mM KCl, 1% Triton X-100, 2 mM DTT, 100 μg/ml cycloheximide and protease inhibitors (EDTA-free, Roche)) and lysates were treated with RNAseI (Ambion) before ultracentrifugation over a sucrose gradient using an SW-41Ti rotor at 222,200 x g for 2 hours at 4 °C. Cytosolic ribosome fractions were collected and digested with PCR grade proteinase K (Roche) in the presence of 1% SDS prior to RNA isolation using TRIzol (Invitrogen). RNA fragments were run on a denaturing urea polyacrylamide gel and correctly sized fragments were eluted in 0.3 M sodium acetate pH 5.5 and recovered by ethanol precipitation. RNA was 3’ dephosphorylated using T4 Polynucleotide Kinase (NEB) and preadenylated linkers were ligated to the RNA fragments using T4 RNA Ligase 2 K277Q (NEB). After 5’ deadenlyase and RecJ exonuclase treatment, fragments were cleaned using an Oligo Clean & Concentrator kit (Zymo) before rRNA depletion by subtractive hybridization using MyOne Straptavidin C1 DynaBeads (Thermo Fischer) and biotinylated rRNA oligos. Following ethanol precipitation, samples were reverse transcribed using SuperScript III RT (Invitrogen) and again ethanol precipitated prior to gel purification using urea PAGE. Fragments were circularized by CircLigase II (Epicentre), recovered by ethanol precipitation, and subjected to library construction PCR using Q5 HF mastermix (NEB), which was size selected using nondenaturing PAGE and recovered by ethanol precipitation. Library quality was confirmed using a Bioanalyzer 2100 (Agilent) with the DNA 7500 kit (Agilent) and libraries were quantified by Qubitt dsDNA BR assay (Thermo Fisher) prior to high-throughput 75 bp single-end sequencing on an Illumina NextSeq machine. Translation efficiencies were calculated from ribosome profiling and RNA sequencing data using Ribodiff version 0.2.1 (52) and differential codon occupancy was determined using diricore as described (39).

### RNA sequencing

RNA was isolated from a portion of cells processed for ribosome profiling using the RNeasy kit (Qiagen). mRNA was isolated by NEXTFLEX® Poly(A) Beads (PerkinElmer), quality checked by RNA 6000 nano chip (Agilent) on a Bioanalyzer 2100 (Agilent) and sequencing libraries were prepared using NEXTFLEX® Rapid Directional qRNA-Seq™ Library Prep Kit (PerkinElmer) according to the manufacturer’s instructions. Library quality was confirmed using a Bioanalyzer 2100 (Agilent) with the DNA 7500 kit (Agilent) and libraries were quantified by Qubitt dsDNA BR assay (ThermoFisher) prior to high-throughput 75 bp paired-end sequencing on an Illumina NextSeq 500 machine. Reads were demultiplexed by sample-specific barcodes and changed to fastq files using bcl2fastq version 1.8.4 (Illumina). An in-house Perl script was used to remove barcodes, and low-quality trimming was performed on both mates using Cutadapt (53) version 1.12 (settings: q=15, O=5, e=0.1, m=20) prior to poly A tail removal using poly A or poly T as an adapter. Reads shorter than 20 bases after trimming were discarded. The trimmed reads were aligned to GRCh38, Ensembl release 95 using HISAT2(54) version 2.1.0 (settings: k=1000, --norc). Reads were only mapped against the forward strand. An in-house Perl script was used to remove reads that mapped to multiple genes. A script from Bio Scientific (dqRNASeq; settings: s=8, q=0, m=1) was used to determine the number of unique fragments per transcript. Unique fragments per gene were determined using an in-house Perl script. Differentially expressed genes were identified using the Bioconductor (https://www.bioconductor.org/) package edgeR(55) version 3.32.1 using R version 4.0.3. Gene expression was normalized using trimmed mean of m-values (TMM), and multiple testing was corrected for using Benjamini-Hochberg correction. Genes with a false discovery rate (FDR) <0.05 and a fold change of at least 1.5-fold were considered differentially expressed. Data were analyzed further using custom R-scripts and topGO version 2.44.0. For gene set enrichment analysis (GSEA), the desktop application (https://www.gsea-msigdb.org/gsea, The Broad Institute) was used to compare the ranked gene list to hallmark signatures using the gene set permutation type with 1000 permutations (36).

### mRNA stability assays

Cells were seeded in 6 cm dishes one day prior to treatment with 1 μg/ml actinomycin D (Sigma-Aldrich) for the indicated time. Cells were washed twice in ice cold PBS and harvested by scraping in ice-cold PBS. RNA was isolated and processed to cDNA as described above for RT-qPCR. The fraction of remaining mRNA after actinomycin D treatment was calculated by the 2^-ΔΔCt^ method.

### MeRIP-qPCR analysis

mRNA was isolated as described above and 1 μg was precipitated with 2 μl anti-m6A antibody (ab151230, Abcam) or normal rabbit IgG (sc-2027, Santa Cruz) in IPP buffer (10 mM Tris-HCl (pH 7.5), 150 mM NaCl, 0.1% NP-40, 1 mM DTT) at 4°C for 4 hours under constant rotation. Then, 20 μl prewashed Dynabeads Protein A (Thermo Fischer) in IPP buffer was added, followed by another 2 hours on the rotator at 4 °C. The beads were washed 5x in IPP buffer and transferred to fresh tubes after the 3^rd^ and 5^th^ washes. Samples were eluted twice in RLT+ buffer (Qiagen) with β-mercaptoethanol for 10 minutes at room temperature and RNA was recovered by ethanol precipitation. Pellets were resuspended in nuclease-free water and subjected to reverse transcription along with 100 ng mRNA input samples followed by RT-qPCR analysis. Transcript recovery over input was calculated and normalized to a positive control mRNA (EEF1A1).

### Statistical analyses

Analysis of ribosome and RNA-sequencing data is described above. For all remaining analyses Student’s t-test or one-, or two-way ANOVA with Dunnett’s post hoc test (when appropriate) was used as described in the figure legends. Statistical testing was performed using GraphPad Prism 6 and statistically significant differences are indicated: *: p<0.05, **: p<0.01, ***: p<0.001.

## Acknowledgements

The authors would like to thank Britt Sterken, Christy Hong and Thamar Jessurun Lobo for experimental advice. This work was supported by a travel grant from the Boehringer Ingelheim Fonds awarded to H.R.Z, and a KWF (#10080) grant awarded to C.F.C. We thank the ERIBA-UMCG sequencing facility for help in preparing the RNA-seq libraries.

## Authorship contributions

H.R.Z. and C.F.C. conceived the project with input from C.M. H.R.Z. performed the experiments with C.M. and G.K. and analyzed the data. F.L-P. provided expertise on the ribosome profiling experiments. R.W. helped with mapping and processing the RNA-sequencing data. E.S. mapped and analyzed the ribosome profiling data. H.R.Z., C.M. and C.F.C. wrote the manuscript. C.F.C. supervised the work.

## Competing interest

The authors declare no competing interests.

**SUPPLEMENTARY Table 1.**
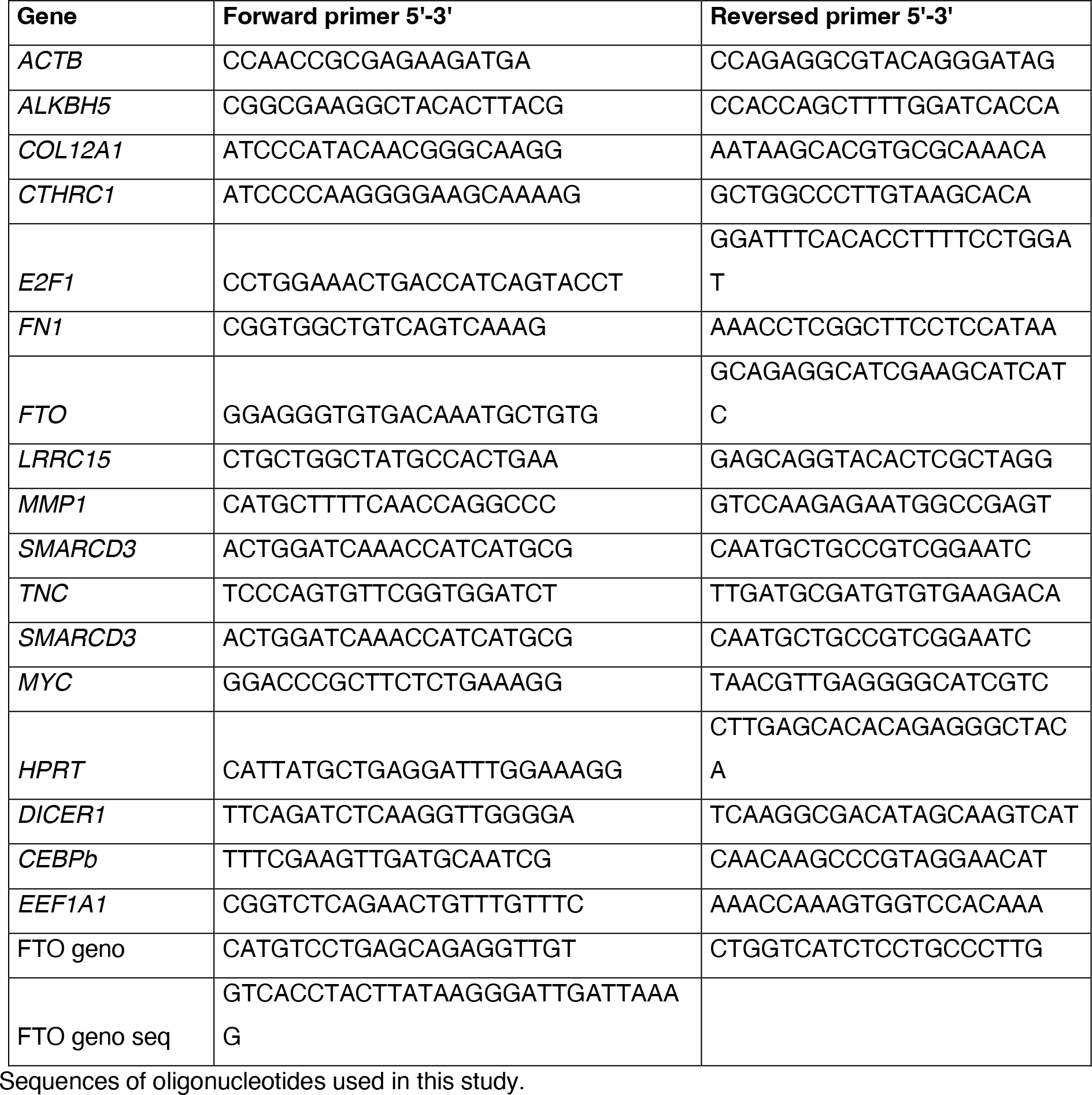

